# Serving as an anatomy laboratory teaching assistant does not improve USMLE Step 1 performance

**DOI:** 10.1101/2025.04.15.648844

**Authors:** Michael A. Pascoe

## Abstract

**Purpose:** Teaching can enhance the teacher’s learning through retrieval practice, elaboration, and metacognitive engagement. In anatomy education, near-peer teaching roles such as laboratory teaching assistants (TAs) may therefore benefit not only learners but also the instructors. This study investigated whether serving as an anatomy TA is associated with improved performance on the United States Medical Licensing Examination (USMLE) Step 1. It was hypothesized that medical students who taught anatomy would demonstrate higher Step 1 scores due to the cognitive benefits of teaching.

**Methods:** Sixty second-year medical students were identified who served as anatomy TAs in a donor-based physical therapy course over seven academic years. Each was matched to a peer from the same graduating class with similar anatomy course performance but no TA experience. De-identified Step 1 scores were obtained from institutional records. An independent samples t-test compared Step 1 scores between groups.

**Results:** The mean Step 1 score was 238.0 (SD = 19.7) for TAs and 236.1 (SD = 16.5) for matched controls. This difference was not statistically significant (t(118) = 0.573, p = 0.568).

**Conclusions:** Medical students who served as anatomy TAs did not perform significantly better on the USMLE Step 1 than their peers. These findings suggest that, under the conditions of this study, participation in a near-peer teaching role did not translate into measurable gains on standardized testing. Educators should consider that while teaching may support professional development and anatomical understanding, additional instructional design elements may be needed to align these experiences with exam-related learning outcomes.

## INTRODUCTION

### Anatomy education in health professions

Anatomical knowledge forms a critical foundation for clinical reasoning, physical examination, and procedural skills across all health professions [1]. Doctor of Physical Therapy (DPT) and Doctor of Medicine (MD) curricula both dedicate substantial instructional time to anatomy in the preclinical phase, often incorporating donor-based laboratory experiences to deepen students’ three-dimensional understanding of human structure [2]. These hands-on laboratory sessions are widely regarded as a pedagogically rich environment for integrating visual, tactile, and spatial learning, and they offer essential opportunities for contextualizing anatomical knowledge with clinical relevance [3]. In both DPT and MD programs, students report that donor dissection enhances their understanding of anatomical variation, strengthens their professional identity, and fosters respectful engagement with the human body as a learning resource [4-6]. However, the resource-intensive nature of donor-based anatomy education requires significant faculty time and laboratory support, which has led many programs to seek instructional innovations that maintain educational quality while addressing scalability [7-8].

### The increasing use of near-peer teaching in anatomy laboratories

One widely adopted solution has been the use of near-peer teaching models, in which students further along in their training provide instruction or support to more junior learners [9]. In anatomy education, near-peer teaching has taken various forms, including structured peer tutoring, supplemental dissection instruction, and formal roles as teaching assistants (TAs) within laboratory settings [10-11]. These models are supported by a growing body of literature indicating benefits for the learners, including increased engagement, reduced anxiety, and improved comprehension of complex spatial relationships [12-13]. Near-peer teaching is also associated with increased relatability, as learners may feel more comfortable asking questions and engaging with instructors closer to their own level of training [14]. In DPT programs, MD students who have already completed their own anatomy coursework often serve as TAs in the anatomy laboratory, providing instructional support and modeling anatomical reasoning [15]. These arrangements not only augment teaching capacity but also create opportunities for interprofessional interaction between future physicians and physical therapists [16]. While the benefits of near-peer teaching for learners are increasingly well documented, much less is known about the educational outcomes for the near-peer instructors themselves.

### Potential benefits to teaching assistants of anatomy

There are strong theoretical and empirical reasons to believe that teaching others may enhance the teacher’s own understanding. Cognitive science has demonstrated that teaching requires active retrieval, elaboration, and metacognitive monitoring, which are processes known to reinforce memory consolidation and deepen conceptual understanding [17-19]. The “protégé effect,” whereby individuals perform better after preparing to teach content to others, supports the idea that teaching can serve as a powerful learning strategy [20]. This may be particularly relevant for medical students preparing for the United States Medical Licensing Examination (USMLE) Step 1, which assesses integration and application of foundational biomedical sciences, including anatomy [21]. The Step 1 exam, developed and administered by the National Board of Medical Examiners (NBME), is a standardized licensure assessment that plays a critical role in a medical student’s educational trajectory [22]. Performance on this exam is a key factor in residency placement and can influence future training opportunities and career direction [23]. As such, Step 1 is widely regarded as a high-stakes milestone in medical education, and considerable effort has been devoted to identifying factors that contribute to strong exam performance

[24-26]. Performance in medical gross anatomy is one such factor that correlates well with Step 1 scores, underscoring the potential importance of strengthening anatomical knowledge through effective study strategies [27-28]. For these students, revisiting and teaching anatomical concepts in a high-fidelity laboratory environment may provide a form of spaced repetition and clinical contextualization that aligns well with evidence-based principles of durable learning [29]. Anecdotal claims and informal faculty observations often suggest that medical students who serve as anatomy TAs perform better on Step 1, particularly in anatomy-related domains, yet no studies to our knowledge have systematically examined this relationship.

This lack of empirical data represents a meaningful gap in the literature on near-peer teaching in health professions education. While prior studies have explored the benefits of peer teaching on learners’ outcomes and on interpersonal skill development for instructors, the potential cognitive and academic benefits, especially in terms of standardized test performance, have received limited attention.

Determining whether serving as a TA reinforces the TA’s own content mastery has important implications not only for optimizing peer-teaching models, but also for informing strategies MD students can use to enhance their own exam preparation. Based on the well-documented benefits of retrieval practice, elaboration, and metacognitive engagement associated with teaching, it was hypothesized that students who served as anatomy TAs would perform better on the USMLE Step 1 compared to their peers. Specifically, it was predicted that there would be a positive association between TA experience and overall Step 1 scores, driven by enhanced consolidation and retrieval of anatomical knowledge.

This study seeks to address this gap by investigating whether second-year MD students who served as teaching assistants in a DPT donor-based anatomy course performed better on the anatomy content of the USMLE Step 1 exam than their peers who did not participate in teaching. In doing so, we aim to evaluate whether the act of teaching contributes to measurable academic outcomes, and to contribute to a growing body of scholarship on the reciprocal benefits of near-peer teaching in anatomy education.

## MATERIALS AND METHODS

### Participants and Group Selection

To examine the influence of participation as a teaching assistant (TA) in anatomy on Step 1 exam performance, we obtained United States Medical Licensing Examination (USMLE) Step 1 scores for all medical student TAs who served in the University of Colorado Physical Therapy (PT) Program over seven consecutive summer terms. Each TA was matched to a control peer from the same medical school graduating class. Control students were selected if their final gross anatomy course grade was within one standard deviation of the corresponding TA, allowing for comparison among students with similar baseline academic performance in anatomy.

### Description of the TA Experience

TAs served during the first summer semester of the PT curriculum, which spanned ten weeks. During this time, two cohorts of PT students were enrolled in either a ten-credit hour course, Clinical Anatomy I (Year 1), or Clinical Anatomy II, a six-week, three-credit hour course (Year 2). Both courses featured whole-body donor dissection and were designed to sequentially cover regional anatomy. Clinical Anatomy I addressed the upper extremity, thorax, head and neck, and lower extremity, while Clinical Anatomy II focused on the abdomen, pelvis, perineum, and joint-specific dissection of the extremities.

Each dissection team consisted of four PT students per donor. Teams followed a combination of commercially available and custom dissection guides [30]. TAs supported the course by facilitating dissection, modeling techniques, responding to student questions, and assisting faculty with lab practical setup outside of scheduled hours. TAs received a brief overview of the experience prior to beginning, although this orientation was not based on best practices, and they were paid hourly out of their work study financial aid funds. Laboratory sessions occurred Monday, Wednesday, and Friday from 1–5 p.m., and TAs reported an average of 12 instructional hours per week.

### Background and Preparation of TA Participants

All medical student TAs had completed their own medical gross anatomy course approximately eight months prior. Their curriculum followed a similar structure with morning lectures and afternoon dissections over a ten-week period and included all regions of the body. The medical anatomy course was graded on a pass/fail basis, and only students who passed were eligible to serve as TAs in the PT course. Medical students volunteered to serve as TAs by expressing interest to the course director a few months prior to the start of the PT anatomy course.

### Data Collection and Processing

TA rosters from the seven cohorts of TAs were submitted to the University of Colorado School of Medicine Data Warehouse Committee. The Committee provided anonymized Step 1 scores for TAs and matched controls, in accordance with institutional data governance protocols. Demographic information was not provided, however, historical enrollment data are available online [31]. Step 1 scores were returned as de-identified datasets and analyzed using Microsoft Excel (Microsoft, Redmond, WA) for organization and initial data cleaning.

### Statistical Analysis

Descriptive statistics were calculated for Step 1 scores in both TA and matched control groups. An independent samples t-test was conducted to evaluate differences in Step 1 performance between TAs and their matched non-TA peers, accounting for the between-class matching design. Statistical significance was set at p < 0.05. All analyses were performed using SPSS (IBM Corp., Armonk, NY).

Prior to conducting inferential analyses, key assumptions of the independent samples t-test were assessed. Shapiro-Wilk tests indicated that Step 1 scores were approximately normally distributed within both the TA group (W = 0.976, p = 0.277) and the control group (W = 0.964, p = 0.072). Levene’s Test for Equality of Variances was not significant (F = 2.556, p = 0.113), supporting the assumption of homogeneity of variance. These results support the appropriateness of the independent samples t-test for comparing group means.

## RESULTS

### Sample Characteristics

A total of 60 medical students served as TAs in the PT anatomy course over the seven consecutive summer terms. The median number of TAs per term was nine and the range was six to 11. Each TA was matched to one medical student peer from the same graduating class, resulting in 60 matched pairs. The only demographic information available revealed that the TA group was a majority male (N = 43; 72%).

### Comparison of Step 1 Scores

To evaluate whether participation as an anatomy TA was associated with performance on the USMLE Step 1 exam, we compared the scores of TAs and matched controls. The mean Step 1 score for the TA group was 238.0 (SD = 19.7), while the control group had a mean score of 236.1 (SD = 16.5).

Results of an independent samples t-test indicated that TAs scored similar to controls, t(118) = 0.573, p = 0.568, suggesting no association between TA participation and Step 1 performance (Figure 1, Table 1).

**Table 1.**
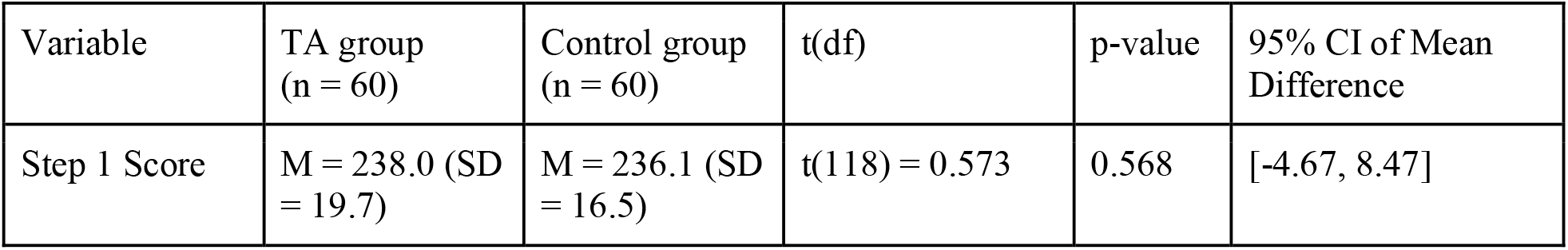
Comparison of Step 1 score between groups. TA = teaching assistant; M = mean; SD = standard deviation; CI = confidence interval.

| Variable | TA group<br>(n = 60) | Control group<br>(n = 60) | t(df) | p-value | 95% CI of Mean<br>Difference |
| --- | --- | --- | --- | --- | --- |
| Step 1 Score | M = 238.0 (SD<br>= 19.7) | M = 236.1 (SD<br>= 16.5) | t(118) = 0.573 | 0.568 | [-4.67, 8.47] |

**Figure 1.**
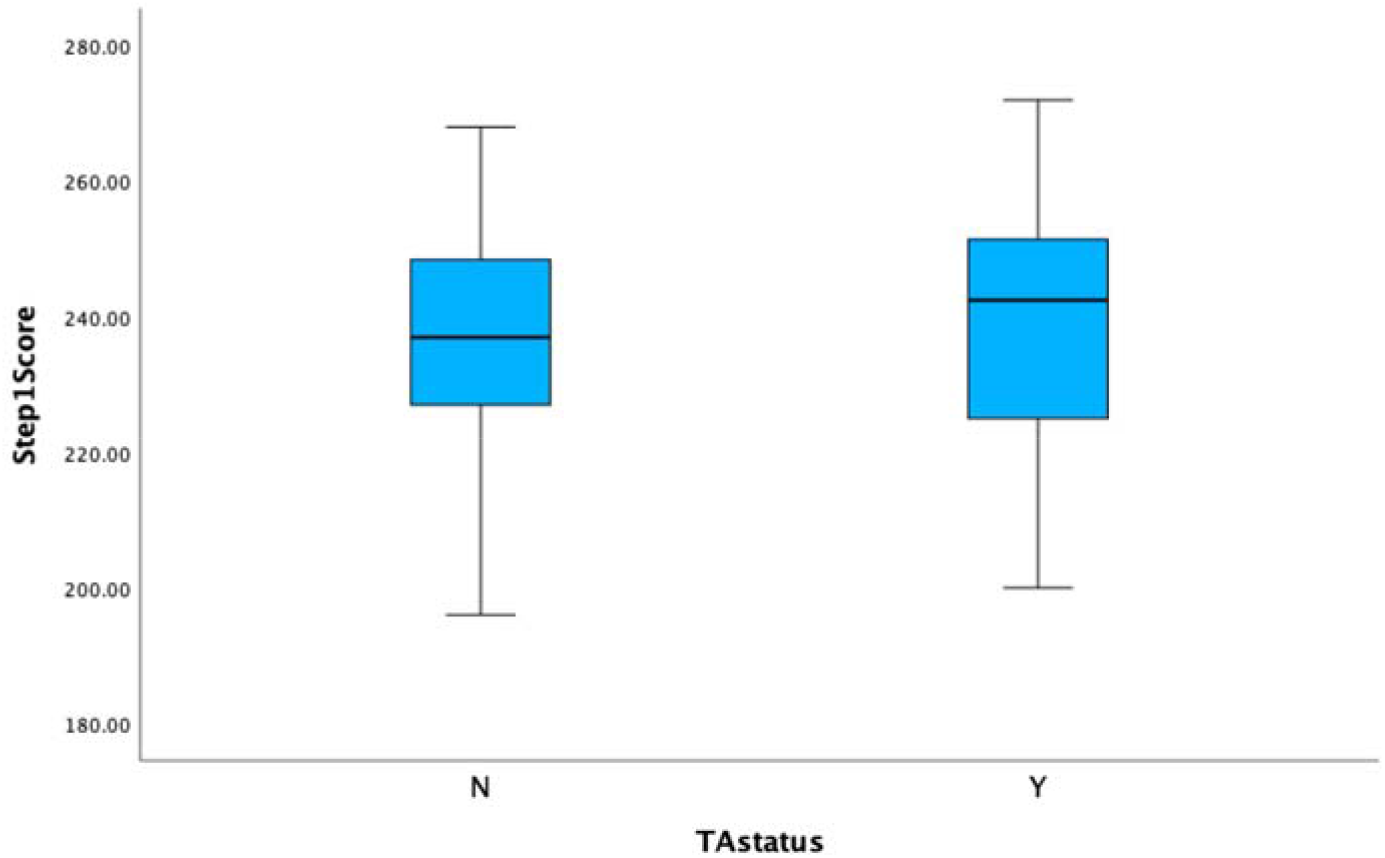
Boxplot of Step 1 scores across both groups.

### Summary of Findings

In summary, medical students who served as anatomy TAs demonstrated no significant difference in Step 1 performance relative to their matched peers. These findings do not support the possibility that participation as a TA in a hands-on, instructional anatomy experience may have educational value with regard to standardized exam outcomes.

## DISCUSSION

### Summary and Interpretation of Findings

This study examined whether serving as a teaching assistant (TA) in a donor-based physical therapy anatomy course was associated with improved performance on the USMLE Step 1 exam among medical students. Despite theoretical and anecdotal support for the cognitive benefits of teaching, no statistically significant difference in Step 1 scores was found between students who served as TAs and matched peers who did not. These results suggest that, within the design and context of this near-peer teaching model, participation as an anatomy TA did not confer a measurable benefit on standardized test performance.

While this finding may be surprising to some educators, it aligns with a growing recognition that the cognitive and academic outcomes of teaching experiences are context-sensitive. Teaching can indeed promote deeper learning through retrieval practice, elaboration, and metacognitive monitoring, as shown in cognitive science literature [17-19]. However, for such benefits to manifest in performance outcomes, particularly on a broad, integrative exam like Step 1, factors such as timing, duration, structure, and intentional preparation may be critical.

### Contextual Considerations and Educational Implications

The TAs in this study completed their own anatomy coursework approximately eight months prior to serving and were not provided with structured pedagogical training. Their teaching activities, while valuable, were relatively limited in weekly hours and may not have aligned closely with Step 1-style question formats or integrative thinking. Moreover, Step 1 assesses a wide breadth of biomedical domains, of which anatomy is only one component (Step 1 Content Outline webpage). Any gains in anatomical reasoning or recall may not have been sufficient to influence total exam scores.

From an educational design perspective, these findings raise important considerations. While near-peer teaching offers clear benefits for learners, such as increased comfort, relevance, and engagement, the benefits for peer instructors may depend heavily on the design of the teaching experience. Programs that aim to promote content mastery through teaching should consider more structured support, including faculty mentorship [32], reflective practice [33], regular feedback [34], and stronger alignment with curricular assessments [35]. These enhancements may help TAs consolidate their knowledge more effectively and transfer it to broader testing contexts [36].

### Limitations and Future Directions

There are several limitations to this study. Matching was based on final anatomy grades, but additional academic or demographic variables were not available due to data anonymization protocols. It is also unknown whether participants who volunteered to be TAs had other distinguishing characteristics, such as higher intrinsic motivation or an interest in teaching, which may confound interpretations of the outcomes [37]. The study was conducted at a single institution with a limited demographic profile, so the generalizability is somewhat limited. Furthermore, this study focused narrowly on Step 1 performance and did not capture other potential benefits of the TA role, including improved confidence, communication skills, or professional identity formation. These outcomes are widely recognized as valuable in medical education [12-14].

Future research should explore not only whether, but how, teaching impacts student learning outcomes. Comparative studies could evaluate the impact of different near-peer teaching models (e.g., structured vs. informal, longitudinal vs. episodic) on both teacher and learner outcomes. Incorporating qualitative data could also reveal how TAs perceive their own learning, and whether these perceptions align with objective outcomes. Such studies could inform more intentional integration of peer teaching within preclinical curricula.

## CONCLUSION

Serving as a TA in an anatomy laboratory did not translate into improved Step 1 performance in this study. While this finding tempers assumptions about the academic benefits of near-peer teaching, it reinforces the need for thoughtful design when engaging students in instructional roles. Educators should remain attentive to the broader professional, interpersonal, and identity-related benefits of these experiences, which may not be captured in standardized test scores but are nonetheless central to the development of future health professionals.

## Acknowledgments

The author acknowledges the University of Colorado School of Medicine Data Warehouse Committee for providing anonymized data that supported this study. A preliminary version of this study was presented in abstract form at the American Association for Anatomy (AAA) Annual Meeting, April 2021.

## DECLARATIONS

### Ethics Approval

The survey and data collection procedures were reviewed and considered exempt by the Colorado Multiple Institutional Review Board (Protocol #18-0514) in accordance with the ethical standards as laid down in the 1964 Declaration of Helsinki.

This manuscript is an original work and has not been published previously. The work is not under consideration in another journal. The author has no relevant financial or non-financial conflicts of interests to disclose. No funding was received for conducting this study. The author has no relevant financial or non-financial interests to disclose. Complete data sets are available upon request.

